# Rheological and Lipid Characterization of Minipig and Human Skin Tissue: A Comparative Study Across Different Locations and Depths

**DOI:** 10.1101/2024.02.25.581953

**Authors:** Harsa Mitra, Evelyn Nonamaker, Ria D. Cordera, Luis Solorio, Arezoo M. Ardekani

## Abstract

Understanding the rheology of minipig and human skin is crucial for enhancing drug delivery methods, particularly for injections. Despite many studies on skin’s viscoelasticity, especially the subcutaneous layer, comparative analyses across different clinical sites are scarce, as is data on the impact of hydration or lipid levels. This study employs shear rheology and lipid analysis to evaluate viscoelasticity and lipid content across three anatomical locations —breast, belly, and neck and three different depth layers in Yucatan minipigs. It reports on how viscoelastic properties change with frequency, time, and strain, noting strain-stiffening and shear-thinning at high strain amplitudes. Human male and female abdominal tissues are also compared to minipig tissues, highlighting distinct viscoelastic traits and lipid’s role in them. The findings suggest the existence of species, anatomical location, tissue depth, and sex-based rheological differences. We also concluded the minipig male tissue is a more accurate model for human male subcutaneous tissue than for females.

## 1 Introduction

Biotherapeutics have emerged as a popular class of drugs due to their potential to treat a wide range of diseases. Among the various routes of administration, subcutaneous (SC) delivery is often preferred due to its convenience, leading to improved patient compliance and reduced healthcare costs. However, the translation of these therapies into clinical use is often hindered by physiological barriers. Specifically, the mechanical properties of the subcutaneous space, including its viscoelasticity, can limit the volume of the drug that can be administered and its subsequent dispersion and absorption. As minipigs provide strong clinical relevance to humans, minipig SC-adipose tissue models have been increasingly used to study pharmacokinetics of various biopharmaceutical injection therapies, especially through the SC route. Therefore, understanding the mechanistic, rheological, and biochemical relationships between minipig models and humans is essential.

The lipid content of the different layers of skin tissue, including the dermis, SC tissue, and hypodermis, can affect the viscoelasticity of the skin, and in turn, the success of SC injections [1, 2]. Higher levels of lipids could result in a more compliant dermis, while lower levels can be associated with a stiffer dermis [3, 4]. This can affect the ease with which a needle can penetrate the skin and deliver the therapeutic drug after the injection. Hence, a stiffer dermis increases the resistance of the skin to needle penetration, affecting drug transport and distribution and potentially leading to injection errors or discomfort for the patient [5]. The needle length and the hypodermis (SC thickness) impact the drug distribution [1, 6, 7]. Additionally, studies have shown correlations between lipid content/water retention and tissue viscoelasticity [2, 8–10]. For example, Lozano *et al.* showed water content contributes to the altered mechanics in collagen-rich soft tissues [8]. Therefore, understanding the viscoelastic and biochemical properties of the skin tissue layers for both the minipig models and humans is important for optimizing the overall performance of autoinjectors and injection therapy and ensuring consistent and effective drug delivery. Although experimental techniques have been previously used to understand biomechanical behavior, the shear rheology research in this area is limited.

In shear rheology, we have two regimes. The first is the linear viscoelastic or LVE regime. Here the tissue deformation is proportional to the applied stress. Consequently, the response is reversible and independent of the magnitude of stress, as long as it’s within a small-strain amplitude range. In contrast, non-linear viscoelasticity occurs outside this range (at large-strain amplitudes), where tissue response is no longer proportional to stress, often irreversible, and dependent on the magnitude and rate of applied stress. More importantly, such non-linearities expose the intra/intercycle processes that the material is undertaking. Malhotra *et al.*, compared the linear viscoelastic moduli under small amplitude oscillatory (SAOS) shear deformations for 3D reconstructed skin models, native-male human, and dermis-only foreskin samples [11]. The study reported that skin elasticity is determined more by the epidermis and body location than by age, directing toward the importance of understanding location-dependent viscoelasticity. A similar comparison was performed by Pan *et al.*, but under large amplitude oscillatory shear deformations (LAOS) [12]. The study found that male human skin exhibited strain stiffening and shear thinning behavior at the highest deformations. We will discuss such shear-dependent intracycle non-linear behavior in detail in Sec. 3.5. The degree of strain-stiffening was reported to be age dependent. Sun *et al.*studied the human abdomen and minipig back adipose tissues under compression and simple shear [13]. They concluded the minipig adipose tissues were stiffer than the human abdominal adipose tissue across their tests [13]. Geerlings *et al.*, also probed the viscoelasticity and described the power-law behavior of the SC-adipose tissue [14]. They suggested that the stiff collagen fiber network that encases the fat globules play a crucial role in the overall mechanical properties of the tissue. These studies provide evidence of the variations in the biomechanical/viscoelastic variations of human cadaver skin at different locations. Therefore, understanding the viscoelastic response of the minipig and human SC tissues from different anatomical locations becomes necessary.

The current study aims to quantify, compare, and describe the dependence of the tissue viscoelasticity based on its total lipid content. We compare three different anatomical locations, i.e., breast, belly, and neck, in a male Yucatan minipig at three different skin-tissue layers. SAOS and LAOS bulk rheology and the Bligh-Dyer method of lipid isolation are used to evaluate the viscoelasticity and lipid content, respectively, across minipig and human skin tissues. An additional case of human belly SC (male and female) is included for comparison purposes. We use the power law and the four-element Maxwell models to extract the various parameters from the frequency and stress relaxation tests. These values can be used for modeling the viscoelastic behavior of various layers at different anatomical locations and will lead to advancing tailored injection techniques for patients. This would improve the overall efficacy of drug delivery through the SC route, minimizing patient discomfort and treatment-related complications.

## 2 Materials & Methods

### 2.1 Tissue Extraction

Tissue collection was conducted in accordance with an approved protocol (# 2011002085) from the Purdue University Institutional Animal Care and Use Committee (IACUC). Male Yucatan minipigs that had been castrated before 3 months of age and weighed between 35 and 40 pounds were used in the experiments. The minipigs were sedated by Purdue University veterinary staff, and the areas of interest (breast, belly, and neck) were shaved. A sterile scrub was then performed, repeated three times with alternating washes of chlorhexidine and sterile gauze. Subsequently, the minipigs were euthanized, and a postmortem scrub was performed, following the same sterile scrub procedure. Tissues from the areas of interest were then extracted using sterile surgical tools and collected in clean plastic bags, which were stored at -80*^◦^*C. Two Caucasian human abdominal SC samples, one male (27-year-old with BMI 26.5) and one female (43-year-old with BMI 31.1), were received from GenoSkin Inc (MA, USA). Both had a Fitzpatrick skin type classification of two. All the tissue samples were placed in clean 2 mL tubes, snap frozen, and stored at -80*^◦^*C.

### 2.2 Rheological Methodology

To probe the viscoelastic behavior of the tissue samples, bulk rheological tests were performed on a Discovery Hybrid Rheometer (DHR) 2 (TA Instruments, DE, USA). The 8 mm stainless-steel sandblasted flat-plate geometry was used along with the Peltier Plate Immersion Cell. The setup is shown in Fig. 1D. The tissue extraction was performed using a 15.87 mm diameter arch punch from the belly, neck, and breast (See Fig. 1A). The rheological properties of tissues may vary with the number of freeze-thaw cycles [15, 16]. Hence, we had employed only one freeze-thaw cycle before the tests were performed. The epidermis, part of the dermis, and muscle layers below the hypodermis were removed with the help of a scalpel. The resulting discs were kept at 2.4 mm thickness (See Fig. 1B). Lastly, to fit the rheometer geometry, 8 mm diameter (2.4 mm thick) discs were extracted using a biopsy punch (See Fig. 1C). Three 8 mm discs were collected from each of the 15.87 mm slices, from the minipig locations, and the male and female human, for triplicate measurements. Once extracted, the samples were allowed to equilibrate at 37*^◦^*C for 15 mins in a petri dish while immersed in 7.4 pH phosphate-buffered saline (PBS) (Gibco, NY, USA). After equilibration, the samples were loaded onto the rheometer geometry and the tests were run at a fixed 2 mm gap (20% compression). Fixing a constant gap height has been shown to produce more consistent results as compared to varying the gap based on the sample thickness [15]. After the sample was loaded and fixed firmly between the plates, 45 mL of PBS 7.4 pH buffer was poured into the immersion cell to avoid sample drying over the period of each test.

**Fig. 1.**
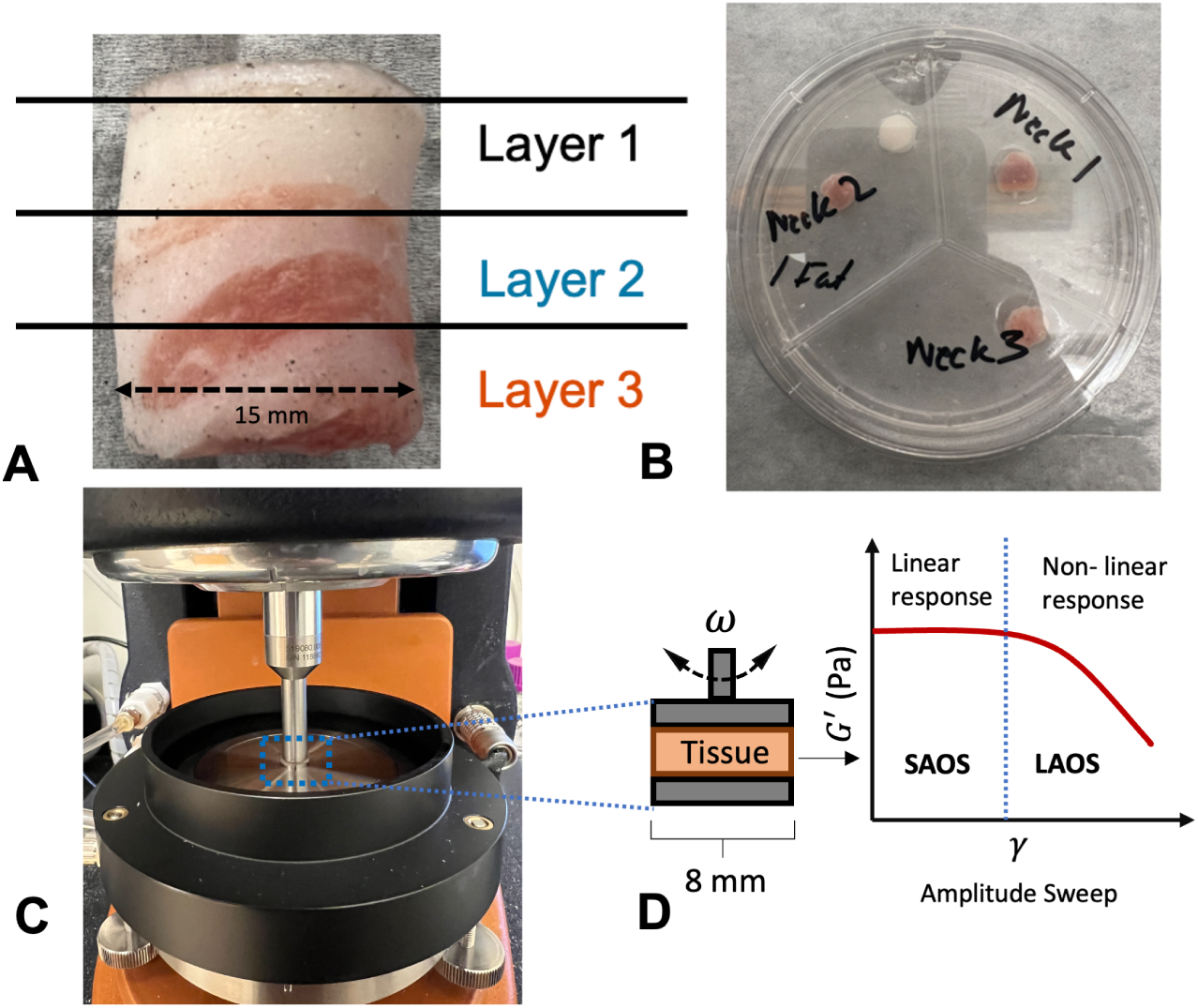
Tissue sample extraction for the rheological tests represented for the minipig neck tissue. A) Punched out tissue with the lines representing the approximated thickness of each layer. Layer 1 excludes the epidermis for all the samples. B) Top view of the sliced layer 1 for the minipig neck tissue. C) DHR-2 Rheometer with the 8 mm parallel-plate geometry and the immersion cell. D) Schematic of the oscillatory shear rheology technique. The amplitude sweep graph schematic shows the difference between the linear (SAOS) and non-linear (LAOS) responses.

To evaluate the rheological properties of the samples, *G^′^* (storage) and *G^′′^* (loss) moduli were measured as a function of frequency and strain amplitude. The frequency sweep at strain amplitude (*γ*) of 0.5% and angular frequency (*ω*) range of 0.04 – 40 rad/s, and amplitude sweep at *ω* =1 and *ω ∈* [0.01–40%] was done. To quantify the non-linear response of the tissues, the stress response waveforms were recorded for each strain amplitude during the strain sweep [15, 16]. All the tests were carried out in a triplicate manner and at 37*^◦^*C while immersed in 45 mL of PBS (pH 7.4).

### 2.3 Lipid Quantification

After rheology testing, the total lipid content of each human and minipig tissue sample was quantified using the following approach based on the Bligh-Dyer method of lipid isolation [17]. It is noteworthy that there may be lipid loss (as the samples were submerged in PBS solution) during the course of the rheological tests due to shearing. Hence, the measured quantities could differ from the initial lipid contents in the samples before each test. Each sample was first dissected into three 20 mg samples. These smaller samples were then homogenized in 200 *µ*L of 8M urea (Sigma Aldrich, MO, USA) using the Qiagen TissueRuptor (Quiagen N.V., Germany). To these tissue homogenates, 800 *µ*L of methanol (Acros Organics, Belgium), 400 *µ*L of chloroform (Sigma Aldrich, MO, USA), and 300 *µ*L of Milli-Q water (Millipore Sigma, MA, USA) were added with vortexing between reagent additions. Samples were then centrifuged at 7000*×*g for five minutes to allow hydrophobic separation to occur. After centrifugation, the upper soluble layer was aspirated, leaving the bottom insoluble layer containing a mixture of lipids and chloroform. The insoluble layer of each sample was then transferred into a clean 0.5 mL tube and placed in the CentriVap Benchtop Vacuum Concentrator (Labconco, MO, USA) at 45*^◦^*C for eight hours to evaporate the remaining chloroform. The total lipid mass for each sample was calculated to be the mass of the dried insoluble pellet. The percent lipid content was calculated to be the ratio of the total lipid mass to the mass of the dissected tissue sample. Statistical analysis between groups was completed using a one-way ANOVA with Tukey comparisons. To visualize tissue composition, histological analysis using a Masson’s trichrome stain was performed on representative cross-sections of tissue from each of the three minipig anatomical locations.

## 3 Results

The mass fractions of the lipid contents are presented along with the rheological evaluation of the minipig and human skin tissues.

### 3.1 Histology and Lipid Contents

Histological examination revealed similar tissue morphologies across the minipig belly and neck cross-sections (Figs. 2A and C). Both tissues contained largely clear striations of the dermis, SC, and muscle tissue, indicated by the regions stained predominantly blue, white, and red, respectively. However, the minipig breast tissue was observed to have larger white adipose deposits throughout, clusters of blue collagen surrounding mammary glands near the center of the tissue, and no clear muscle layers (Fig. 2B).

**Fig. 2.**
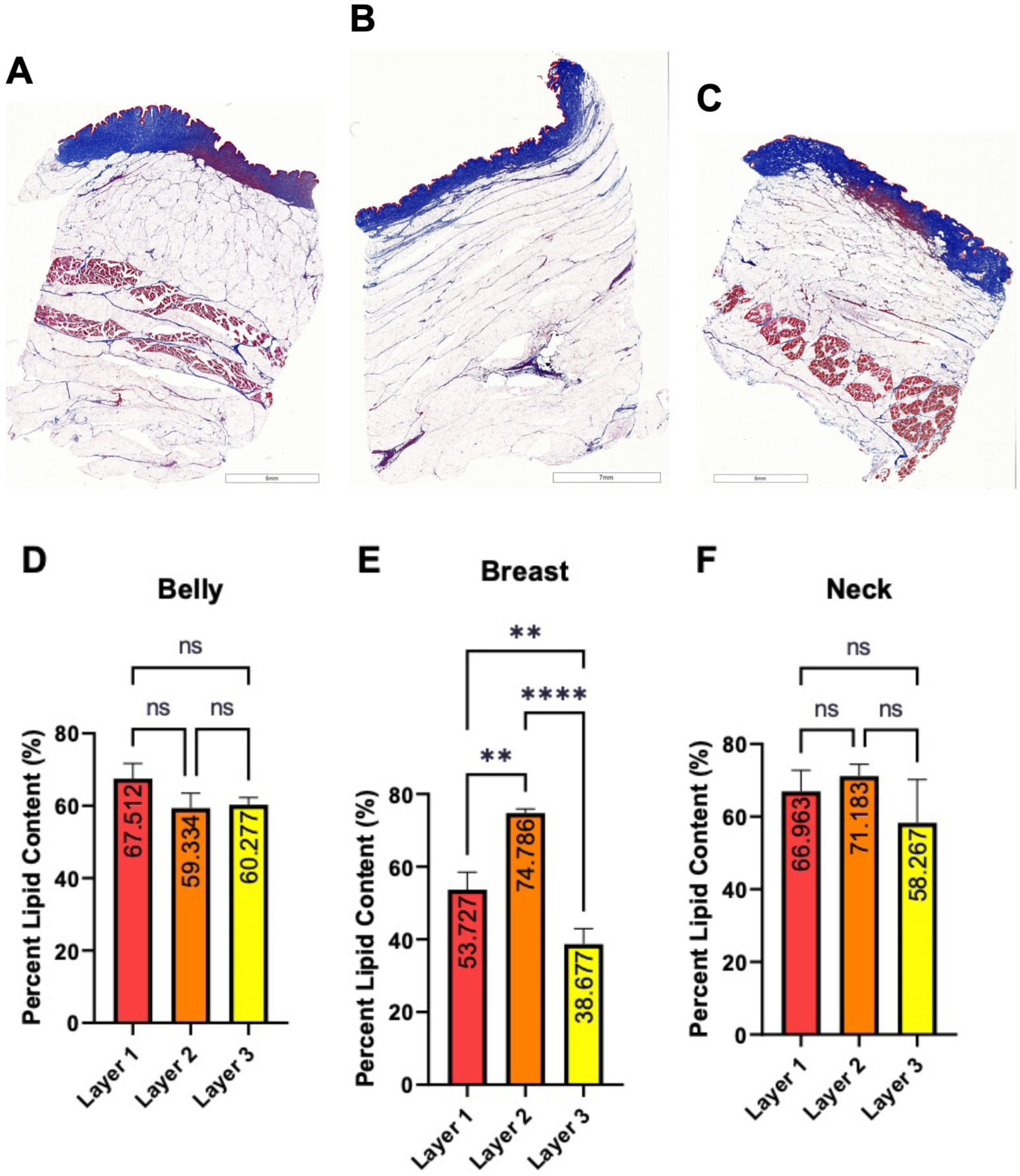
Comparison of percent lipid content across minipig anatomical locations. A) Masson’s trichrome stain of minipig belly tissue. B) Masson’s trichrome stain of minipig breast tissue. C) Masson’s trichrome stain of minipig neck tissue. D) Percent lipid content of tissue layers obtained from the minipig belly region, post-rheology testing. E) Percent lipid content of tissue layers obtained from the minipig breast region, post-rheology testing. F) Percent lipid content of tissue layers obtained from the minipig neck region, post-rheology testing. Data in D-F are reported as the mean ± SD of nine 20 mg samples resulting in no significance (ns) or p-values (*∗∗* = *p ≤* 0.01, *∗ ∗ ∗∗* = *p ≤* 0.0001).

The minipig belly and neck tissues also behaved similarly regarding their lipid quantification. In both tissues, the total lipid content remained relatively constant across each of the three tissue depths, at about 60-65% of the total sample mass after rheological testing (Figs. 2D and F). On the other hand, lipid analysis of the minipig breast samples indicated significantly different quantities of lipid across tissue depths. Layer 2, corresponding to the middle 2 mm of the original tissue biopsy punch, had the highest total lipid content at approximately 75% of the sample mass post-rheology testing whereas layers 1 and 3 contained only 54% and 39% lipid mass, respectively (Fig. 2E).

Upon analyzing the lipid content of the human samples, it was discovered that the female samples contained about 20% more lipids than the male samples (Fig. 3A). Both the male human and female data were compared to the minipig data to determine which minipig anatomical region and depth best correlated to the human tissues in terms of total lipid content. The lipid content of the male human samples was most similar to that of the minipig belly and neck samples across tissue depths, with the exception of layer 2 of the minipig neck (Figs. 3A, C, and D). On the other hand, the lipid content of the human female samples was most similar to the minipig breast and neck tissues in layer 2 with marked statistical differences when compared to all other minipig tissues at all other depths (Figs. 3B, C, and D).

**Fig. 3.**
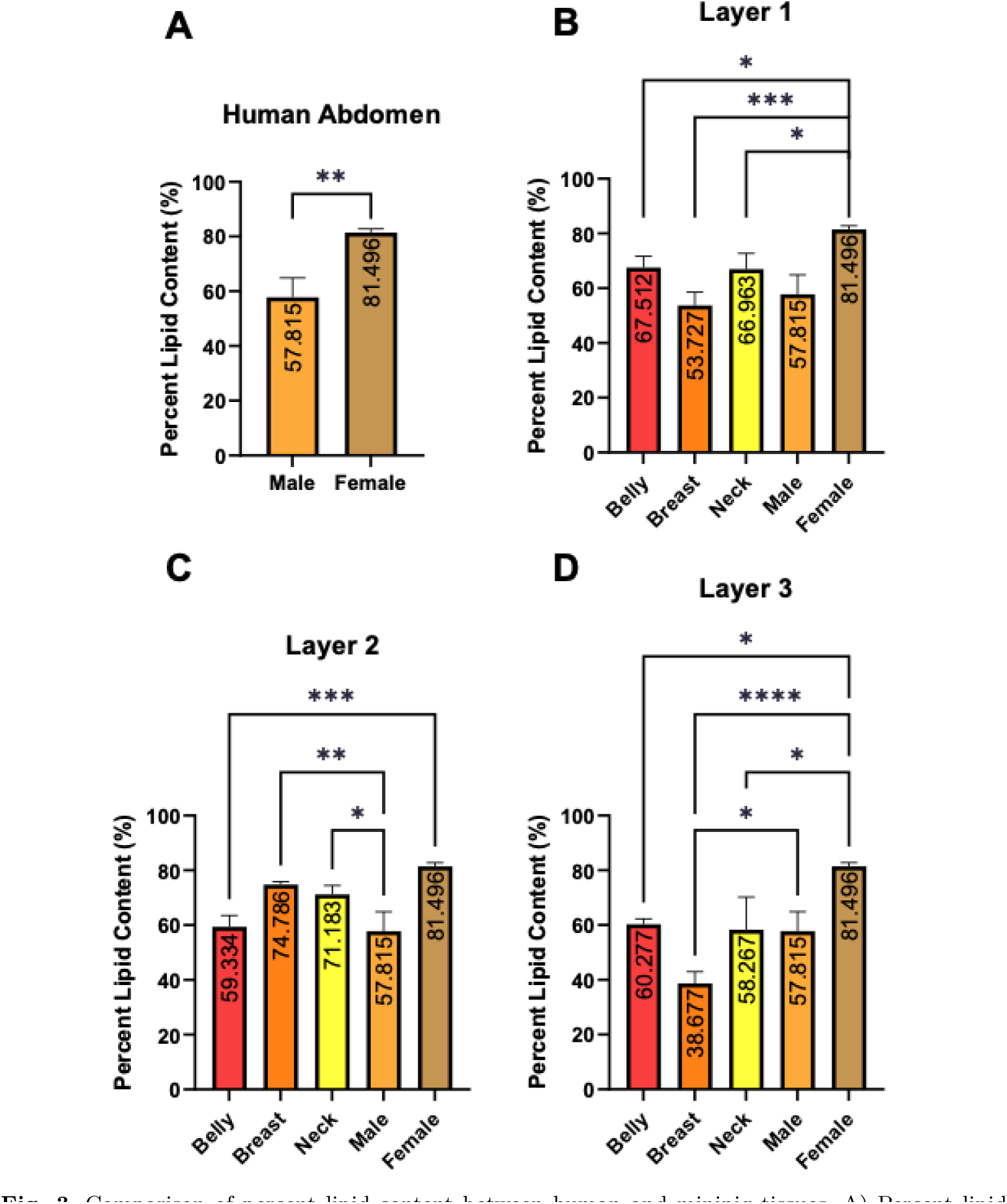
Comparison of percent lipid content between human and minipig tissues. A) Percent lipid content of human tissue samples isolated from GenoSkin models. B) Comparison of percent lipid content between human samples and layer 1 of all minipig anatomical locations. C) Comparison of percent lipid content between human samples and layer 2 of all minipig anatomical locations. D) Comparison of percent lipid content between human samples and layer 3 of all minipig anatomical locations. Data in B-D are reported as the mean ± SD of nine 20 mg samples resulting in no significance (ns) or p-values (*∗* = *p ≤* 0.05, *∗∗* = *p ≤* 0.01, *∗ ∗ ∗* = *p ≤* 0.001, *∗ ∗ ∗∗* = *p ≤* 0.0001).

### 3.2 Frequency Sweep Tests

The frequency sweep results are presented in Figs. 4 and 11. The applied strain amplitude for all the tests was 0.5% (SAOS). Thus, the response is expected to be within the linear viscoelastic (LVE) regime. Across all frequency sweeps, the storage modulus exhibits power-law behavior, with the storage modulus being higher than the loss modulus for all cases. Among all the tested tissues, the minipig neck layer 2 and breast layer 1 showed the highest and lowest storage modulus, respectively. Considerable differences were observed for both the, *G^′^* and *G^′′^*, in Figs. 4 and 11. In Fig. 4a, i.e., layer 1, the neck region (breast region) shows the highest (lowest) storage modulus. A similar trend, i.e., *G^′^* : *Neck > Belly > Breast*, except for, *ω >* 5 *rad.s^−^*^1^, can be observed for the case of layer 3 in Fig. 4c. Although in Fig. 7b, we observed, i.e., *G^′^* : *Neck > Breast > Belly*. The two human SC samples exhibited close agreement to the minipig belly layer 2 in Fig. 4b. Also, it should be noted, in Fig. 4a, the human SC samples behaved stiffer, i.e., had a higher storage moduli, as compared to the Minipig- belly and -breast layer 1. Additionally, all the minipig layer 3 samples had higher storage moduli than the human SC samples (Fig. 4c).

**Fig. 4.**
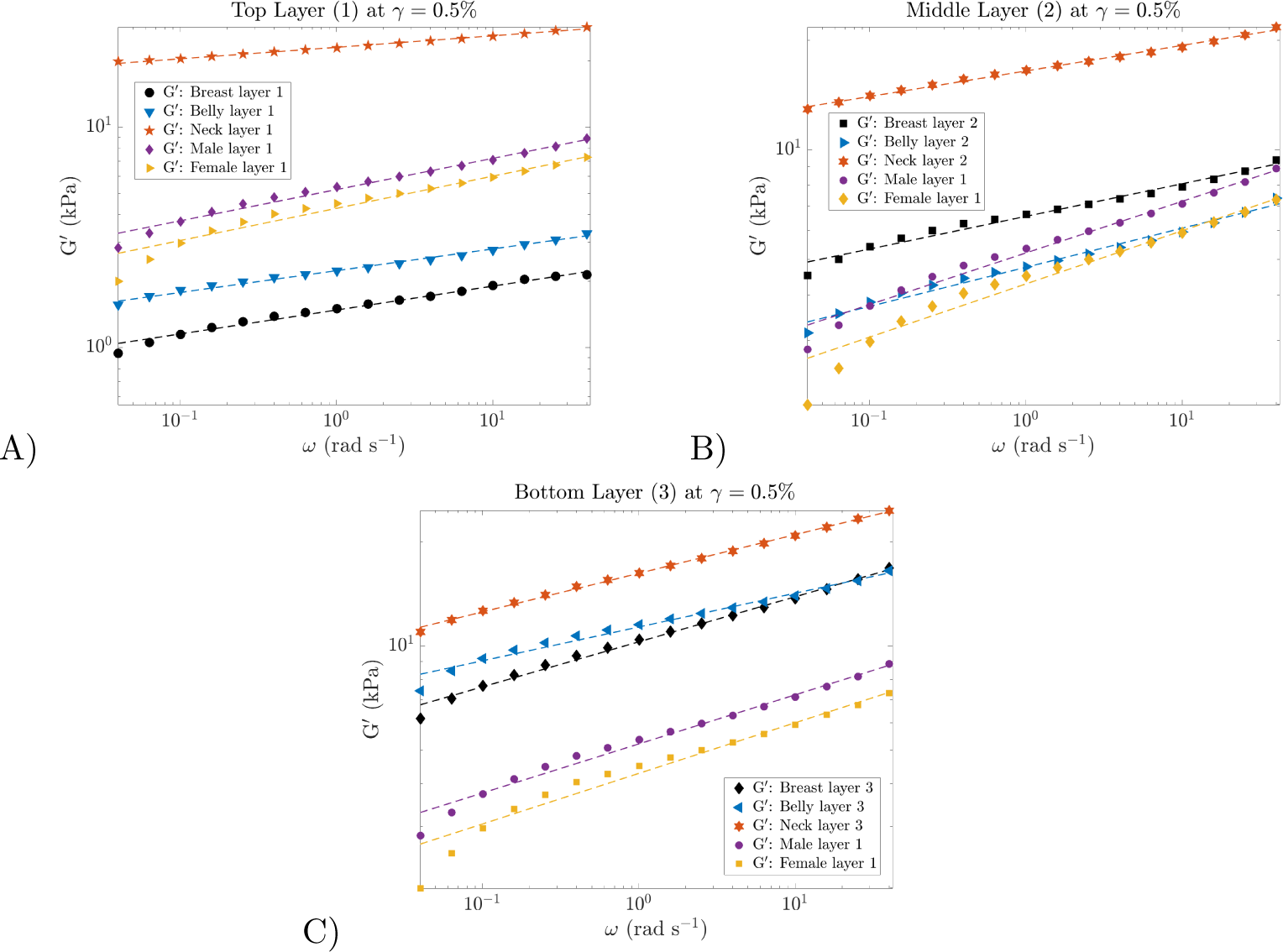
Storage modulus, G*^′^*(*ω*), for the three different regions of the minipig, i.e., breast, belly, and neck, across the different layers, 1-3, represented through a-c, respectively. Dashed lines present the power law fit. The male and female human SC samples are overlaid in all the figures for comparison.

In Figs. 11A-C, the loss modulus for all the minipig layers and locations, along with the male human and -female SC layers, have been presented. The female SC showed a lower storage and loss modulus as compared to the male counterpart. Tissues that exhibit higher lipid content (here the female human SC) are recognized for their low storage modulus. This phenomenon can be attributed to the comparatively lower inter-molecular forces and greater flexibility of lipids, as compared to other biomolecules such as proteins and carbohydrates [18]. Therefore, tissue layers containing a greater quantity of lipids are likely to demonstrate increased flow and deformation when subjected to shear stress, resulting in a lower storage modulus. Although, it is important to note that other factors, such as the degree of saturation of the lipids, also play a role in overall storage and loss modulus [19, 20]. A trend, i.e., *G^′′^* : *Neck > Belly > Breast*, similar to the elastic counterpart is observed for the loss modulus across the minipig layer 1 and 2 (Figs. 11A and B, respectively).

In terms of the lipid contents in the minipig, belly, breast, and belly possessed the highest fractions in layers 1, 2, and 3, respectively (See Figs. 3B, C, and, D), though the neck showed the highest storage modulus in all three layers in Fig. 4. When *G^′^* of the belly and breast are compared in Fig. 4, we observe the belly, breast, and belly to have the highest storage modulus, especially at lower frequencies. As previously explained, the tissues with higher lipid content, tend to show higher viscous behavior. Despite of that, the female SC has the highest lipid content as compared to the male SC (See Fig. 3). It exhibited lower loss moduli as compared to the male SC, as observed in Fig. 11. Similar deviations can be observed in Fig. 4A, where *G^′^*(*ω*) for the minipig neck is higher as compared to its belly. Although, the lipid fraction in the minipig belly and neck were quantified to be fairly similar (See Fig. 3B). These observations go on to show challenges with performing shear rheology of high lipid content, i.e., soft tissues [2]. Although, with the appropriate selection of the rheometer geometry, sample preparation and loading, and temperature control, such challenges can be minimized.

A power-law relation is fitted to the storage moduli across Figs. 4a-c, based on,

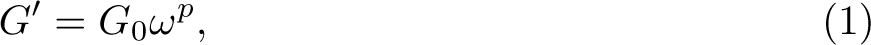

where *G*_0_, and, *p*, are constants [14, 21, 22]. The fitted parameters are represented in Fig. 5 and Table 1. In Fig. 5, it can be observed the material with the lowest (highest) *G^′^*(1) had the highest (lowest) power index, *p*. The fitted power-law model with R-square values greater than 0.985 for all the tested cases (Refer to Table. 1).

**Fig. 5.**
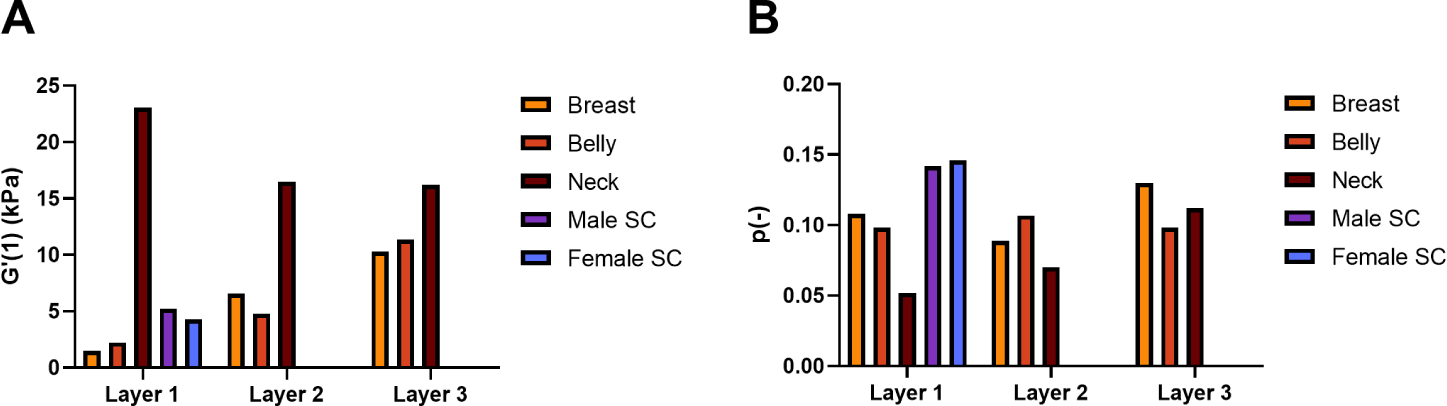
Visualization of the fitted parameters for the frequency sweep data in Table 1. A) *G^′^*(1), and B) *p*, for all the layers and locations are displayed. The male human/female SC fits are represented with the minipig layer 1 for both parameters for comparison.

**Table 1.** Power-law fit for the storage modulus, *G^′^*(*ω*), for the various locations and layers

|  | Layer 1 | Layer 2 | Layer 3 |
| --- | --- | --- | --- |
| **Minipig Breast** |  |  |  |
| $G'(1)$ (kPa) | 1.480 | 6.565 | 10.280 |
| $p$ (-) | 0.108 | 0.089 | 0.130 |
| **Minipig Belly** |  |  |  |
| $G'(1)$ (kPa) | 2.232 | 4.758 | 11.340 |
| $p$ (-) | 0.098 | 0.107 | 0.098 |
| **Minipig Neck** |  |  |  |
| $G'(1)$ (kPa) | 23.050 | 16.470 | 16.21 |
| $p$ (-) | 0.052 | 0.070 | 0.112 |
| **HM Belly** |  |  |  |
| $G'(1)$ (kPa) | 5.210 | - | - |
| $p$ (-) | 0.142 | - | - |
| **HF Belly** |  |  |  |
| $G'(1)$ (kPa) | 4.283 | - | - |
| $p$ (-) | 0.146 | - | - |

**Fig. 6.**
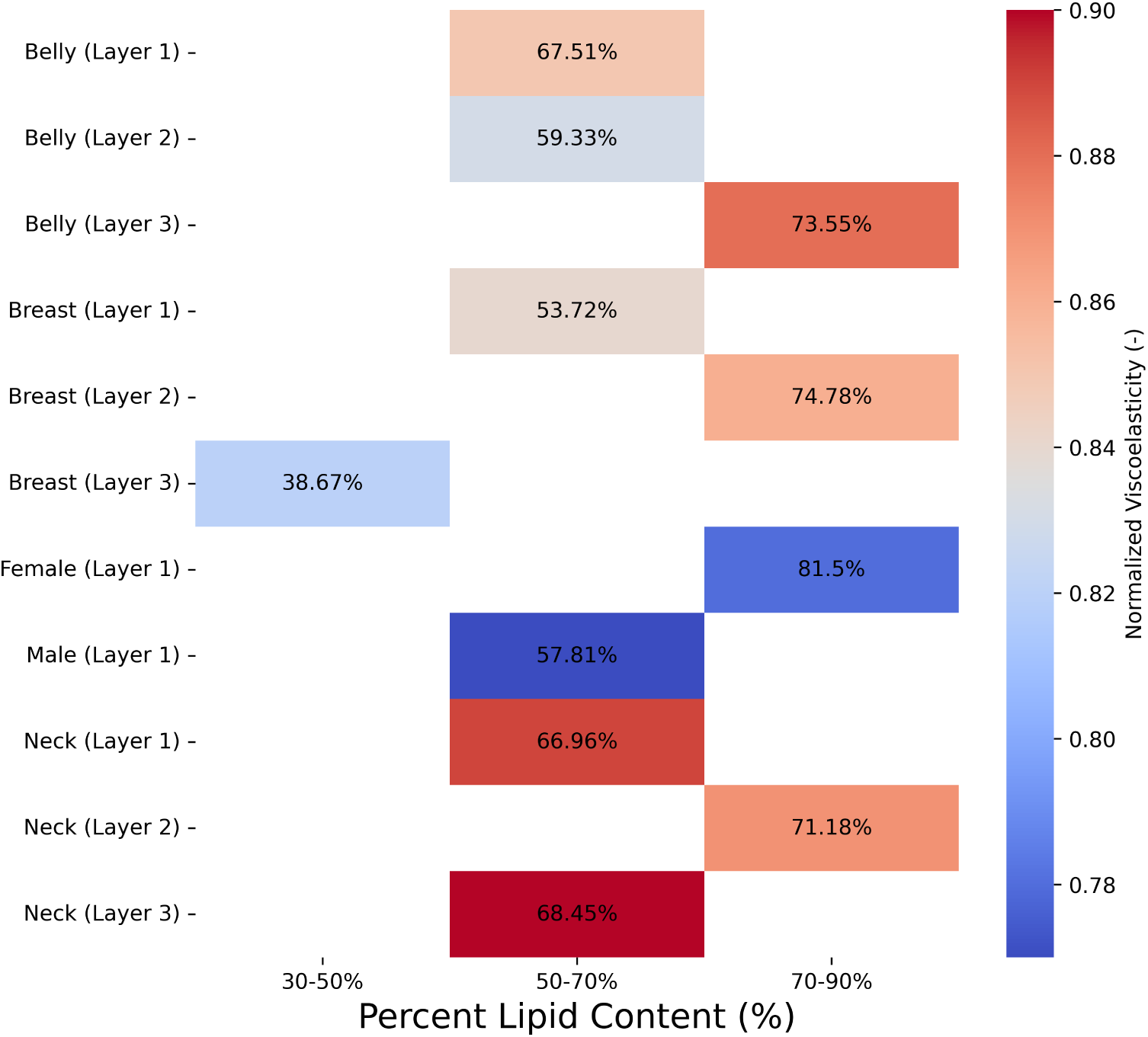
A colored heat map of the normalized viscoelasticity at *ω* = 1 rad/s, i.e., purely viscous/elastic at a value of zero/one, w.r.t. percent lipid content. Additionally, each viscoelastic value for the respective tissue is overlaid with the corresponding percent lipid content. The female and male layer 1 represents the human SCs. A correlation of, *r* = 0.73*, p* = 0.024, is observed among all the minipig tissue viscoelasticity and the lipid content.

### 3.3 Amplitude Sweep Tests

The averaged storage and loss modulus values from the triplicate tests for all the tissue samples are presented in Figs. 7 and 12, respectively. An average variation of 16.5% was observed between the three triplicates. When compared among each minipig anatomical location, the breast and belly regions showed strong similarities across their *G^′^*(*γ*), and *G^′′^*(*γ*), values among each layer, especially at lower strain amplitudes.

Unlike the layers of minipig belly and breast tissue, the three layers from the neck had a narrower range of moduli. Similar to the frequency sweep results for human tissues presented in Sec. 3.2, the female SC (highest lipid content) showed lower storage and loss moduli at lower strain amplitudes as compared to the male counterpart (See Figs. 7 and 12). Also, the storage modulus of both the male human and female SC layers presented a similar behavior to the minipig-belly and breast layer 2, as observed in Fig. 7B. Specific to the storage modulus, as seen in Fig. 7, layer 1 compared to layers 2 and 3, showed earlier onset of non-linear viscoelastic response [15, 16, 23, 24]. Interestingly, based on the lipid quantification, each layer did not have any significant difference. But it should also be noted, in the case of the minipig belly, despite having a similar lipid content (Refer to Fig. 2D) in each layer, *G^′^*(*γ*) and *G^′′^*(*γ*) had strong variations across each layer (See Figs. 7 and 12). Comparing the magnitude of *G^′^*(*γ*) across the three minipig layers in Fig. 7, we observe a similar trend to the frequency-sweep results, as expected and as explained in Sec. 3.2. Based on the evaluations shown in Figs. 7 and 12, the minipig-neck layer 2 and layer 3 showed the highest magnitude of storage and loss modulus, respectively, across *γ* = 0.01 to 40%, among all the test cases. The minipig-breast layer 1 showed the lowest-magnitude moduli. For the SC layers (layer 1), the minipig-breast region had lower moduli as compared to the belly (See Fig. 7A and Fig. 12A. Compared to the minipig-belly and breast SC tissues, the male human and female SC tissues had higher moduli. (See Fig. 7A and Fig. 12A).

**Fig. 7.**
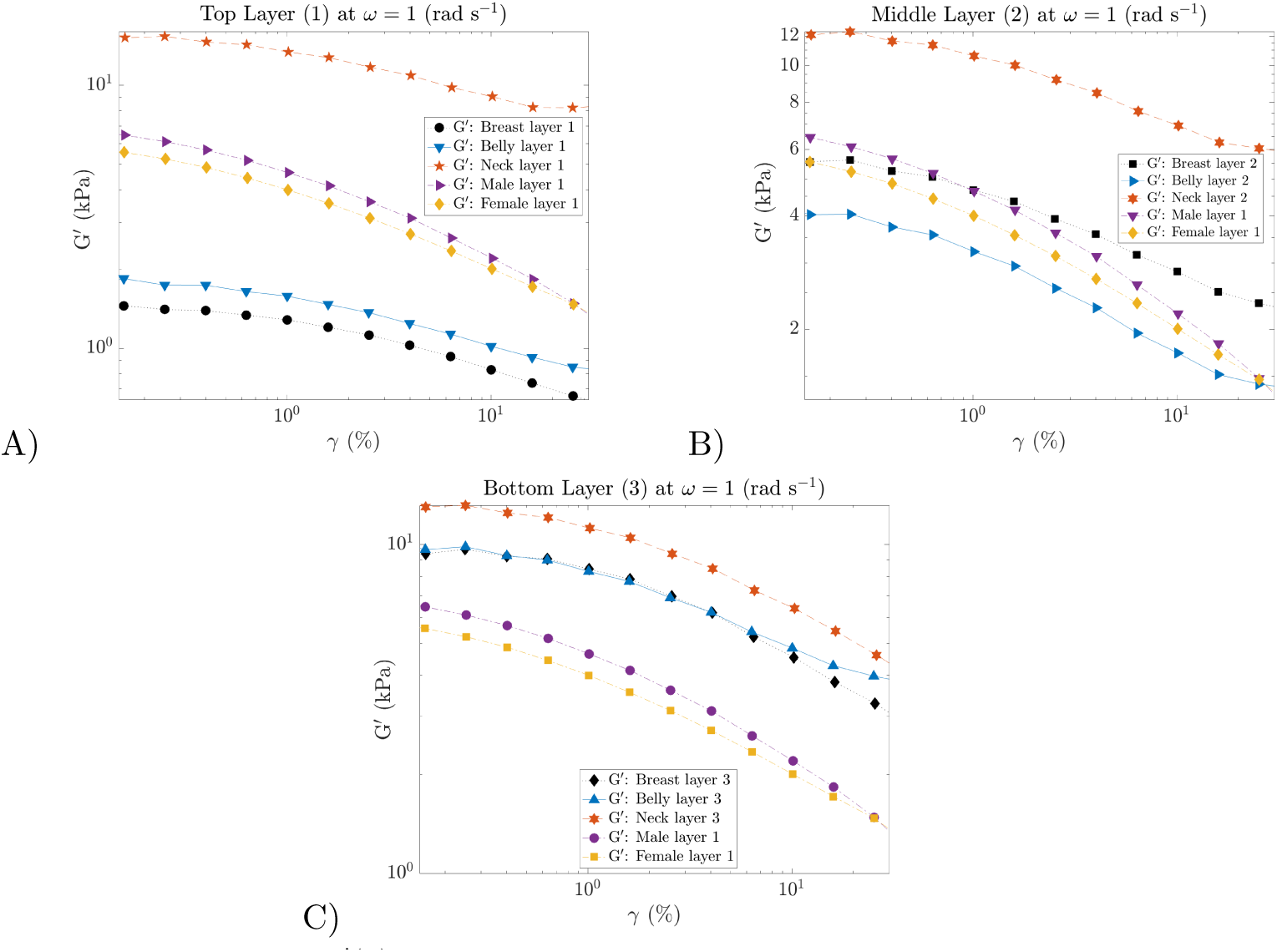
Storage modulus, G*^′^*(*γ*), for the three different regions of the minipig, i.e., breast, belly, and neck, across the different layers, 1-3, represented through A-C, respectively. The male and female human SC samples are overlaid in all the figures for comparison. Solid lines are overlaid to guide the reader and are not a fit.

Additionally, within the linear region, storage and loss moduli are independent of the applied strain amplitude (at a constant frequency). This results in a sinusoidal wave. In the nonlinear region, storage and loss moduli are a function of the strain amplitude when the frequency is kept constant. Consequently making the stress waveforms distorted from sinusoidal waves [25]. We will discuss the non-linear response in detail in Sec. 3.5. Using this information, across Figs. 7 and 12, among the lower two layers, i.e., 2 and 3, layer 2 had an earlier onset of nonlinearity. A higher degree of non-linearity, especially at higher strain amplitudes, is observed in the human SC tissues, as compared to the minipig cases in Fig. 7 and Fig. 12. In brief, close agreement between the minipig belly and breast were observed, especially for layers 1 and 3. Layer 3 showed the highest similarity for all the minipig tissue cases. The minipig-neck layers showed similar storage modulus across the various layers, although the behavior is reduced at higher-strain amplitudes.

### 3.4 Stress Relaxation Tests

The relaxation behavior of the tissue samples under a constant application of strain is quantified and represented in Fig. 8. All the tests are performed at a constant shear strain of 40%, specifically probing the non-linear region. Across the triplicate tests, the magnitude of *G*(*t*) across the three tests, varied at an average of 11%. The test results, *G*(*t*), are fit to the four-element generalized-Maxwell (GM), also known as Maxwell–Wiechert model, and can be described by the following equation [26],

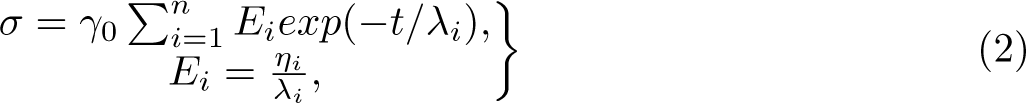

**Fig. 8.**
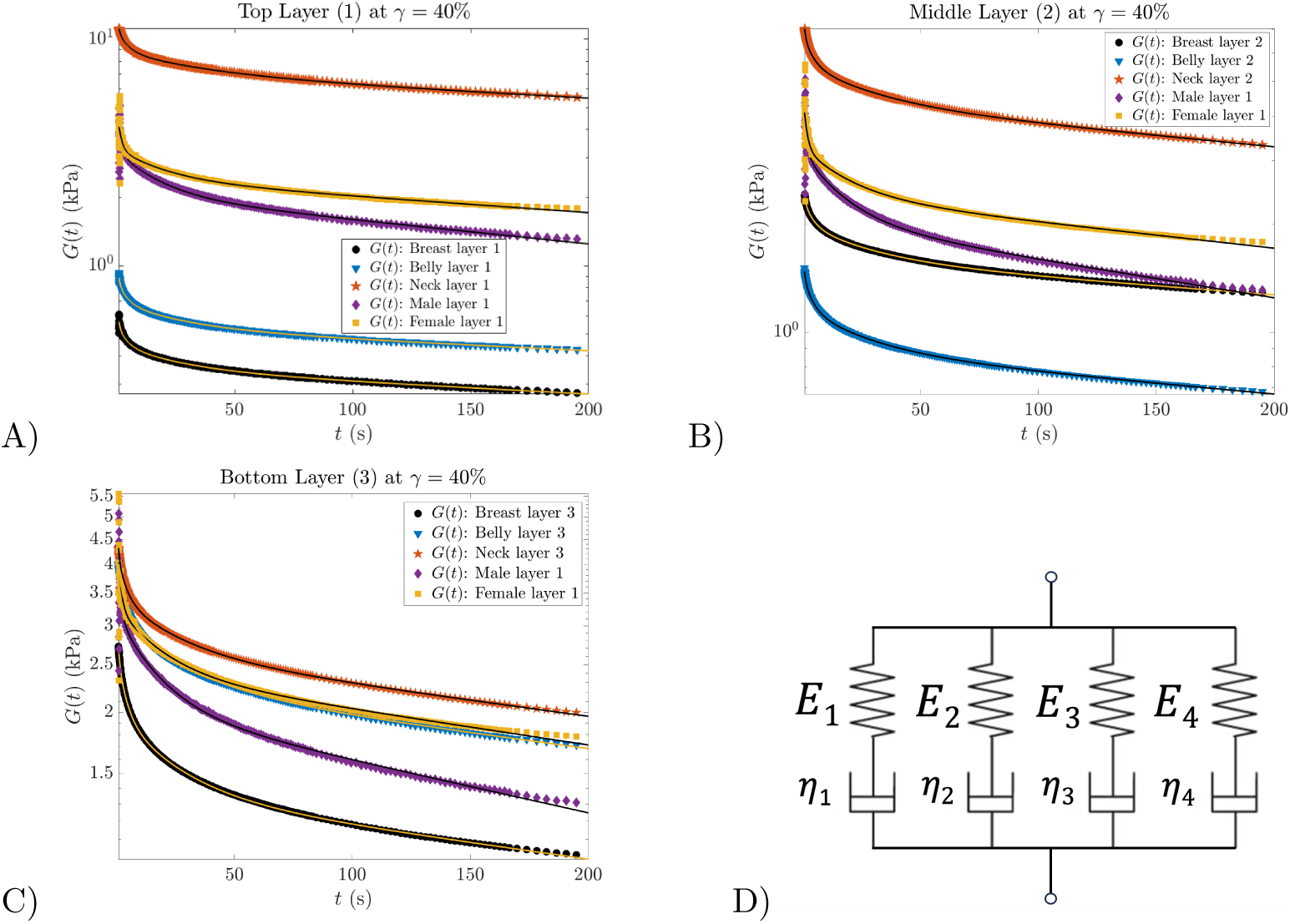
Relaxation modulus, G(t), for the three different regions of the minipig, i.e., breast, belly, and neck, across the different layers, 1-3, represented through a-c, respectively. The male and female human SC samples are overlaid in all the figures for comparison. Solid lines represent the four-element GM-model fit. The four-element GM model is represented in D. Each arm corresponds to one Maxwell element.

where, *σ*, *γ*_0_, *η_i_*, and *λ_i_*, are the total stress response, constant applied strain at *t* = 0, viscosity, and relaxation time of the of the *i^th^* Maxwell element, respectively. In this study, we used a four-element GM model, i.e., *n* = 4, which showed the best fit for all the relaxation data (See Fig. 8D). The fitted parameters are tabulated in Table 2. For better comparison, the maximum and minimum elastic modulus and relaxation time, i.e., among the four elements, are presented in Figs. 9A and B, respectively. The above GM model parameters can be utilized to calculate the frequency-dependent storage and loss moduli as follows [26],

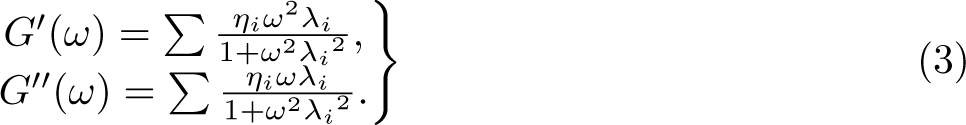

**Fig. 9.**
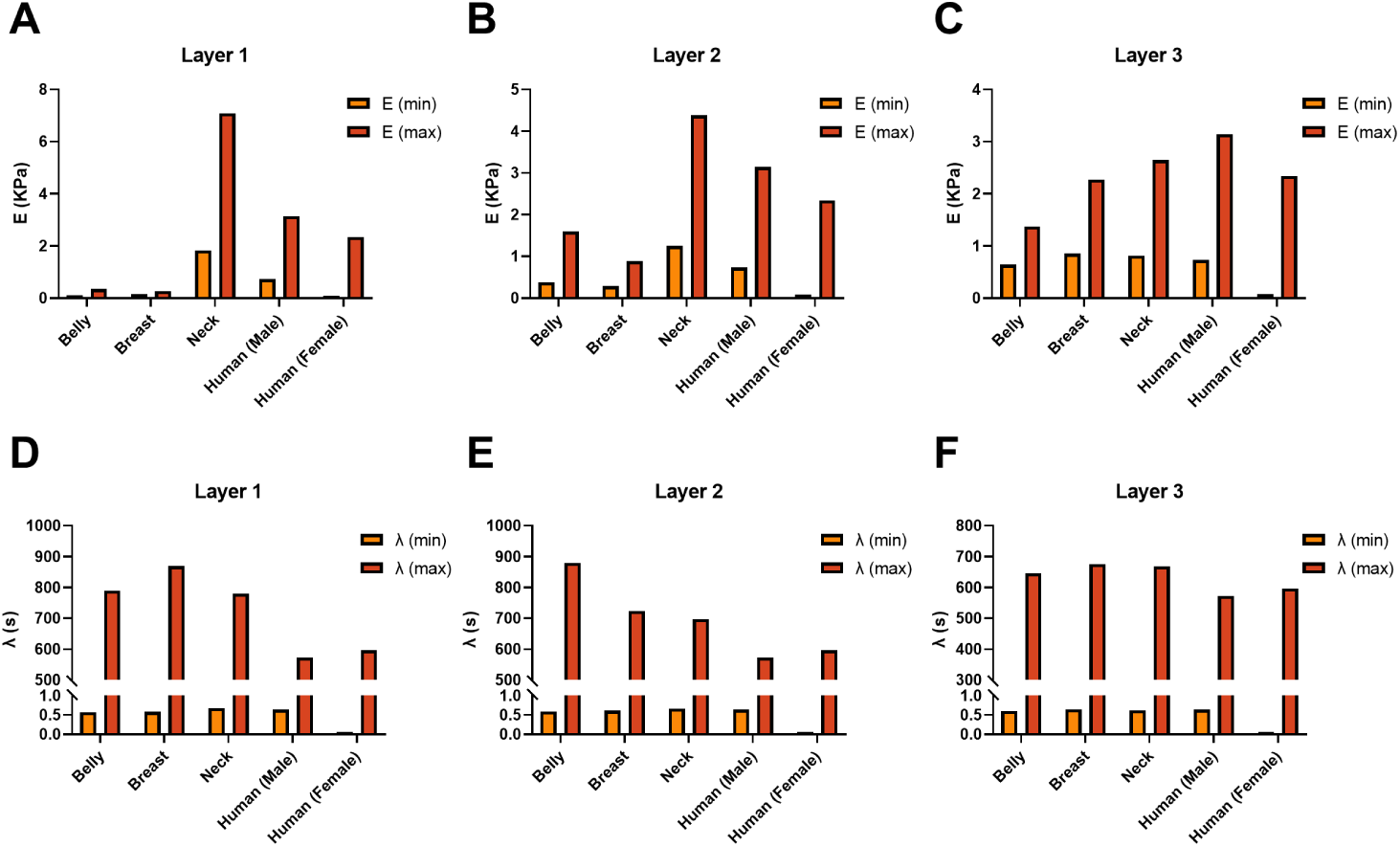
GM-model fit parameters of *G*(*t*). The maximum and minimum of the four elements, A) elastic modulus, *E_i_*, and B) relaxation time, *λ_i_*, of all the various layers for the minipig samples, i.e., top (Layer 1), middle (Layer 2), and bottom (Layer 3). The human, i.e., male and female, data is only for the SC (top layer) but is overlaid in all the figures for comparison purposes. It is important to note, that the minimum relaxation time values are negligible as compared to their respective maximums. Hence are not visible in b. Table 2 can be referred to for exact values of, *E_i_*, and, *λ_i_*, for all the samples tested.

**Table 2.** Four-element Maxwell model fit on the stress relaxation behavior, *G*(*t*), for all the three anatomical locations and depth layers.

|  | Layer 1 | Layer 2 | Layer 3 |
| --- | --- | --- | --- |
| **Breast** |  |  |  |
| $E_1$ (kPa) | 0.1039 | 0.4224 | 0.9387 |
| $E_2$ (kPa) | 0.1628 | 1.596 | 0.7168 |
| $E_3$ (kPa) | 0.1033 | 0.3774 | 1.3680 |
| $E_4$ (kPa) | 0.3496 | 0.4596 | 0.6396 |
| $\lambda_1$ (s) | 3.50 | 28.61 | 0.60 |
| $\lambda_2$ (s) | 0.57 | 879.80 | 3.64 |
| $\lambda_3$ (s) | 25.39 | 3.68 | 645.70 |
| $\lambda_4$ (s) | 789 | 0.59 | 25.00 |
| **Belly** |  |  |  |
| $E_1$ (kPa) | 0.1639 | 0.8862 | 2.2660 |
| $E_2$ (kPa) | 0.2603 | 0.2843 | 1.1120 |
| $E_3$ (kPa) | 0.1747 | 0.3788 | 0.8563 |
| $E_4$ (kPa) | 0.5304 | 0.2863 | 0.8576 |
| $\lambda_1$ (s) | 27.07 | 723.60 | 675.70 |
| $\lambda_2$ (s) | 0.58 | 3.86 | 0.65 |
| $\lambda_3$ (s) | 3.62 | 0.61 | 4.02 |
| $\lambda_4$ (s) | 869.10 | 27.93 | 27.93 |
| **Neck** |  |  |  |
| $E_1$ (kPa) | 2.0690 | 1.2580 | 0.8165 |
| $E_2$ (kPa) | 2.2320 | 4.3800 | 0.8492 |
| $E_3$ (kPa) | 7.0860 | 1.6400 | 1.1910 |
| $E_4$ (kPa) | 1.8330 | 1.2490 | 2.6470 |
| $\lambda_1$ (s) | 4.25 | 29.89 | 25.78 |
| $\lambda_2$ (s) | 0.68 | 697.10 | 3.59 |
| $\lambda_3$ (s) | 779.70 | 0.66 | 0.62 |
| $\lambda_4$ (s) | 35.52 | 4.01 | 667.80 |
| **HM Belly** |  |  |  |
| $E_1$ (kPa) | 0.7321 | - | - |
| $E_2$ (kPa) | 1.822 | - | - |
| $E_3$ (kPa) | 0.9219 | - | - |
| $E_4$ (kPa) | 3.144 | - | - |
| $\lambda_1$ (s) | 6.157 | - | - |
| $\lambda_2$ (s) | 572.8 | - | - |
| $\lambda_3$ (s) | 34.68 | - | - |
| $\lambda_4$ (s) | 0.6434 | - | - |
| **HF Belly** |  |  |  |
| $E_1$ (kPa) | 1.585 | - | - |
| $E_2$ (kPa) | 0.0782 | - | - |
| $E_3$ (kPa) | 2.399 | - | - |
| $E_4$ (kPa) | 0.8892 | - | - |
| $\lambda_1$ (s) | 1.417 | - | - |
| $\lambda_2$ (s) | 0.0046 | - | - |
| $\lambda_3$ (s) | 596.5 | - | - |
| $\lambda_4$ (s) | 20.12 | - | - |

Additionally, the relaxation half-life times [27], *λ*_1*/*2_, i.e., timescale at which the stress is relaxed to half its initial value, is presented in Table 3. A clear difference between the half-times is observed between the minipig and human tissues, with the former exhibiting longer half-lives. A shorter half-life time results in a faster remodeling of the tissue matrix under stress. Moreover, we notice, the neck region’s half-life times are relatively similar across the layers, suggesting a more uniform relaxation behavior. The breast location showed the most variability in their half-life times among each layer. In Fig. 8A and C, the magnitude of the relaxation modulus follows the order of *G*(*t*) : *Neck > Belly > Breast*. Although, between each location, the difference in the magnitude of *G*(*t*) is more significant in layer 1 (See Fig. 8A), especially between the neck and belly/breast. In Fig. 8B, the minipig-layer 2, the relaxation modulus magnitude followed the trend of *G*(*t*) : *Neck > Breast > Belly*. Additionally, the human SC samples had a closer agreement with the minipig breast layer 2 (See Fig. 8B). Noticeably, the female SC showed the strongest correlation to the belly layer 3, as observed in Fig. 8C. The human female SC layer had a higher degree of relaxation as compared to the other male human SC and minipig layers.

**Table 3.** Relaxation half-life times, *λ*_1*/*2_, for all the three anatomical locations and depth layers.

| | $\lambda_{1/2}$ (s) | | |
| --- | --- | --- | --- |
|  | Layer 1 | Layer 2 | Layer 3 |
| Breast | 48.7 | 186 | 25.4 |
| Belly | 71.8 | 84.9 | 57.1 |
| Neck | 165 | 120.5 | 164.5 |
| Male | 22.8 | - | - |
| Female | 55.1 | - | - |

As discussed in previous sections (See Sec. 3.2 and 3.3), tissues/tissue layers with higher lipid content will have a lower storage modulus. This is evident across the male human and female SC in Fig. 9A. Although, in Fig. 9B, the maximum relaxation time for both the male human and female SC tissues were similar, suggesting that a faster stress-relaxation behavior is not necessarily accompanied by a higher viscous behavior. Therefore, it is important to note that the relationship between lipid content and stress relaxation can be more complex and could depend on other factors not quantified here.Even though the neck has the highest elastic modulus (See Figs. 9A.i, ii, and iii). As seen in Figs. 9B. i and ii, the belly and breast, in layers 1 and 2 showed the longest relaxation times. All the minipig layers showed similar relaxation times in layer 3 (See Figs. 9B.iii). In brief, the relaxation behavior of the tissues followed a four-element GM model. In comparison to all the tissue cases, the male human SC showed the fastest relaxation. This can be observed from the fast relaxation times for the male and female human SC in Figs. 9B.i, ii, and iii.

### 3.5 LAOS Tests

The recorded strain waveforms at each strain amplitude during the amplitude sweep tests were post-processed using MITLaos [28]. The shear stress response can be described as a sum of higher harmonic contributions as [28, 29],

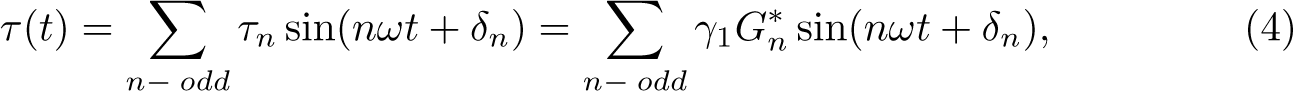

where *G^∗^*, *γ*_1_, and *δ_n_* are the complex moduli, maximum applied strain, and the phase angle, respectively. Using the framework described by Ewoldt *et al.*, intracycle non-linearities, strain stiffening/softening, and shear-thinning/thickening behavior can be quantified using the following set of variables [29]. The periodic stress response at a steady state is plotted parametrically against the strain response. Such representations are called Lissajous–Bowditch (LB) plots. Material properties can then be determined graphically using the LB plots. In which *τ*, i.e., total-shear stress, is plotted as a function of *γ* for the elastic behavior. Whereas the same is plotted as a function of **γ̇** for the viscous behavior. The minimum strain elastic shear modulus or tangent modulus is evaluated at *γ* = 0, *G^′^_M_*. While the large strain elastic shear modulus is quantified at the maximum imposed strain (*γ* = *γ*_1_), *G^′^_L_*. This framework can be understood using the following parameters for a sinusoidal stress/strain input,

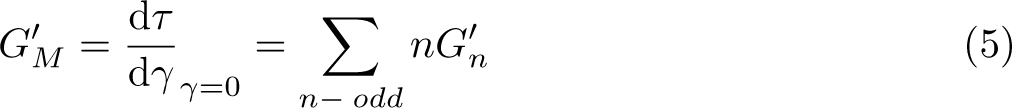

and,

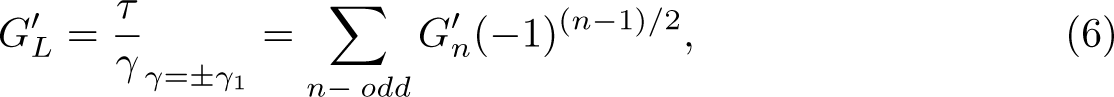

Similar properties can be quantified for viscous components using the Fourier parameters of the higher harmonic stress contributions. The minimum-rate and large-rate dynamic viscosities, *η^′^_M_* and *η^′^_L_*, respectively, can be defined as following[28, 29],

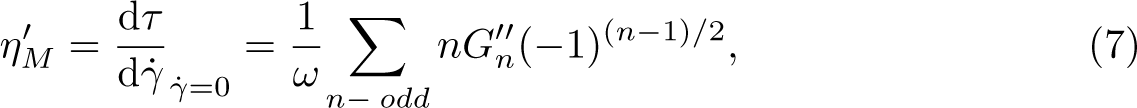

and

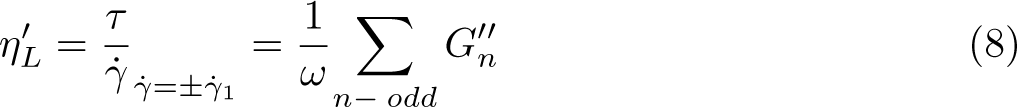

where **γ̇** is the shear-rate. Based on these sets of variables, a strain-stiffening ratio is defined as[29]:

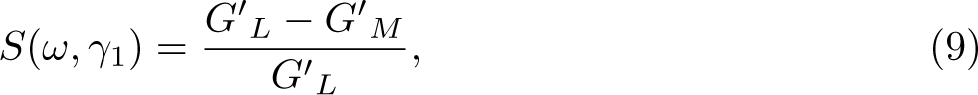

and the shear-thickening ratio:

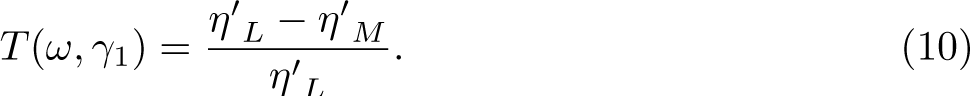

For cases when S *>* 0, the elastic behavior of the material can be interpreted as intra-cycle strain-stiffening, whereas S *<* 0 indicates intra-cycle strain-softening, S = 0 for a linear elastic response. The S parameter becomes crucial in understanding tissue functionality, as it relates to how tissues maintain structural integrity or adapt under physiological stresses. Similar to the elastic behavior, the viscous counterpart, T *>* 0 corresponds to intra-cycle shear-thickening and T *<* 0 indicates intra-cycle shear-thinning. Also, it is to be noted that, T = 0 corresponds to a linear viscous response[29]. In simpler terms, the T parameter describes the tissue’s ability to change under deformation at different shear rates. This is vital to quantify tissue fluidity and its response to dynamic loading. In summary, a high positive S value combined with a higher magnitude of T (when T*<* 0), results in a higher degree of stiffening as strain is applied. But also accompanies a faster dissipation of the developed non-linear elastic stresses through the shear thinning behavior of the tissue. The S and T parameters are quantified for all the tissue samples at strain amplitudes of 15 and 40%. All the results shown are at *n* = 3 and *ω* = 1 rad. s*^−^*^1^.

For brevity, only the S and T values for *γ*=15% and 40%, are visualized in Fig. 10 and also tabulated in Tables 4 and 5. Additionally, the LB plots are presented for all the tissue cases in Fig. 13. All of the tissues from both the minipig and human skin showed strain stiffening and shear thinning behavior. This can be observed from the S and T values in Tables. 4 and 5. It should be noted the non-linear behavior is sensitive to the testing conditions, such as compression, pre-straining, etc. Previous observations of strain softening and strain thinning in tissues have been also reported [12, 30, 31]. As compared to the human SC layers, higher values of S and T are observed for the minipig tissue layers, especially at the higher strain amplitude of 40%. Except for the elastic behavior of the female SC. In conclusion, all the tissue cases showed strain stiffening and shear thinning behavior. The female-SC tissue with the highest lipid content (81.49%) showed the lowest degree of strain-stiffening behavior with, *S* =0.03 and *S* =0.15, for 40% and 15% strain, respectively. Accompanied by an increased strain amplitude. However, the female SC showed an increase in the shear-thinning degree, compared to the rest of the tissue cases. This can be explained by the negative correlation between lipid content and stiffness [3]. Apart from the female SC, all the tissues showed a higher degree of strain-stiffening behavior at higher strain amplitude (See Table 4 and 5).

**Fig. 10.**
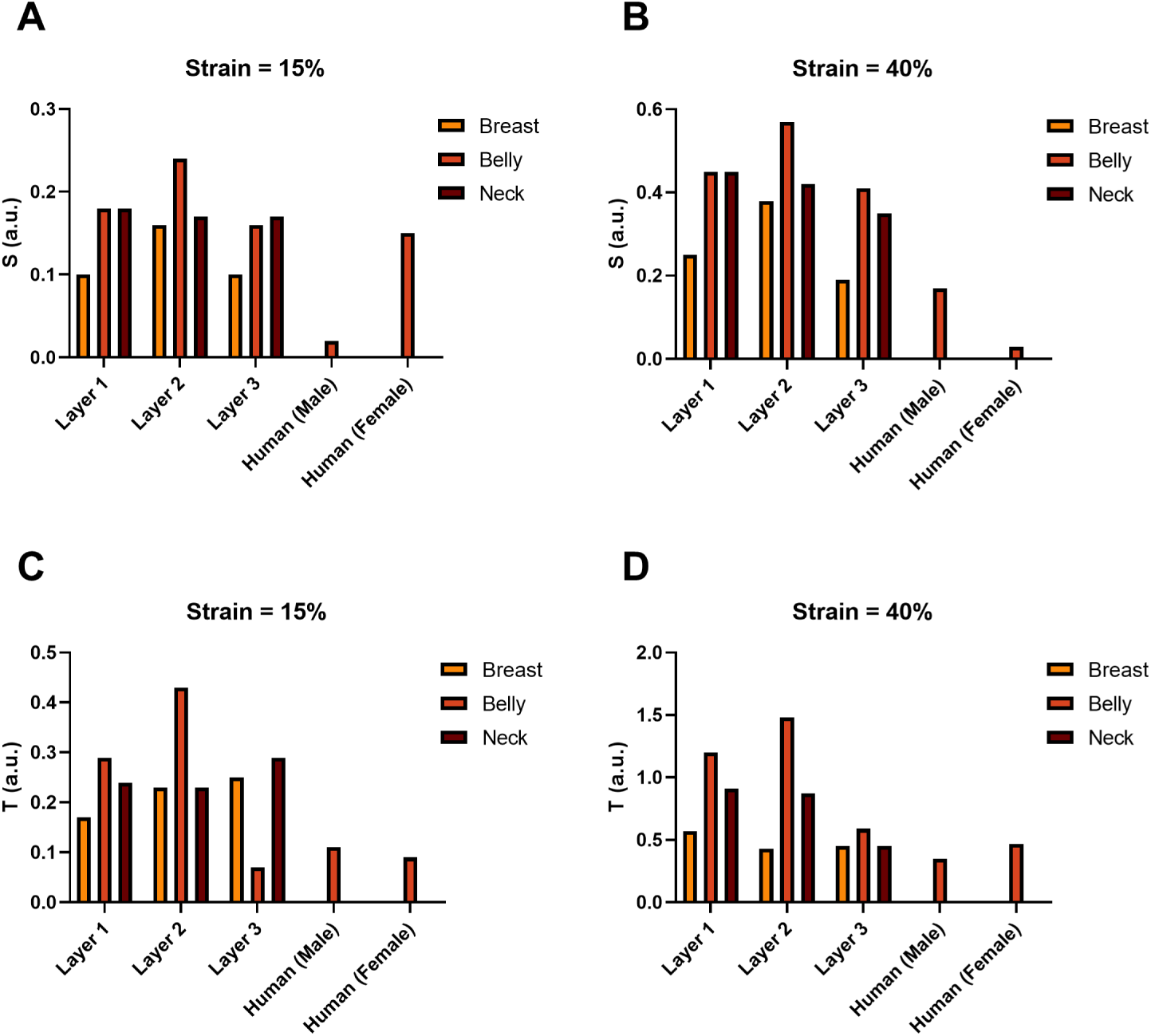
A) and B) represent the S (elastic behavior) values for both the strain conditions of 15%. and 40%, respectively. C) and D) show the T (viscous behavior) values for 15% and 40% applied strain. The respective S and T values are tabulated in Tables 4 and 5.

**Table 4.** S and T parameters at *γ*_1_ = 15% for all the samples tested. HM/HF Belly represents male human/Female Belly, respectively.

|  | Layer 1 (SC) |  | Layer 2 |  | Layer 3 |  |
| --- | --- | --- | --- | --- | --- | --- |
|  | S | T | S | T | S | T |
| **Breast** | 0.10 | -0.17 | 0.16 | -0.23 | 0.10 | -0.25 |
| **Belly** | 0.18 | -0.29 | 0.24 | -0.43 | 0.16 | -0.07 |
| **Neck** | 0.18 | -0.24 | 0.17 | -0.23 | 0.17 | -0.29 |
| **HM Belly** | 0.02 | -0.11 | - | - | - | - |
| **HF Belly** | 0.15 | -0.09 | - | - | - | - |

**Table 5.**
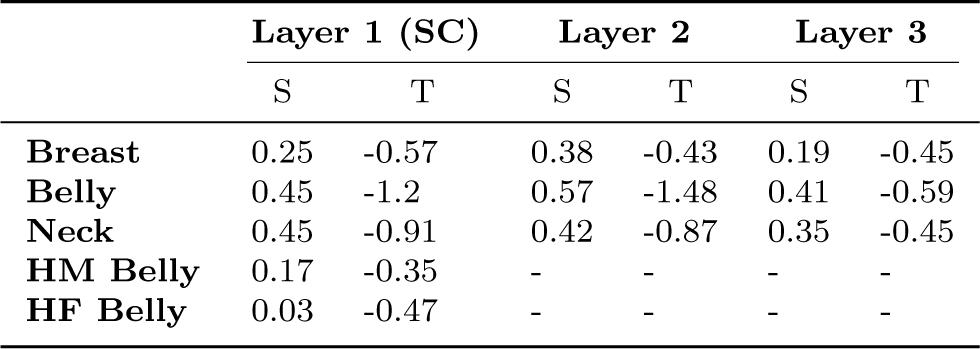
S and T parameters at *γ*_1_ = 40% for all the samples tested. HM/HF Belly represents male human/Female Belly, respectively.

## 4 Discussion

The biochemical composition of tissue, namely its lipid content, plays a role in driving its viscoelastic properties via hydrophobic interactions. To explore this hypothesis, we extracted the total lipid content of minipig tissue samples isolated from the belly, breast, and neck region of castrated male Yucatan minipigs and human tissue samples isolated from the commercially available GenoSkin model. The total lipid fractions were accompanied by the differences in tissue viscoelasticity across the different locations, depth layers, and the two species.

The total lipid content across tissue depths and species was compared to see whether similarities in lipid content would be observed across samples that behaved similarly during mechanical testing. In examining the total lipid content of minipig samples, we discovered the distribution of lipids remained relatively constant across layers in both the belly and neck tissues. On the other hand, significant differences in lipid content were observed across tissue layers in the minipig breast tissue, with the largest proportion occurring in the layer corresponding to the middle 2 mm of the original biopsy punch. These measurements correlated with histological findings illustrating that the neck and belly tissues share similar morphological characteristics, whereas the breast tissue contains a higher proportion of adipose and glandular deposits, particularly in the middle region.

We observed that the human (female) samples had a higher proportion of total lipids compared to the male human samples. This agrees with prior findings indicating human female SC tissue tends to have higher numbers of adipocytes when compared to its male counterpart [32]. This, combined with our earlier observations in the minipig breast tissue, suggests biological sex and the presence of high levels of estrogen could be factors contributing to the recruitment of cell phenotypes that drive the accumulation of lipid material. When the lipid content of the minipig and human tissue samples were compared, the highest degree of similarity was observed between the minipig belly and neck samples and the male human samples. On the other hand, the lipid content of the human female samples was significantly different from nearly all minipig samples, with the exception of Layer 2 of the minipig breast and neck tissues. These groupings were also observed in the stress relaxation response (See Sec. 3.4) of the tissue samples. Taken altogether, these findings suggest the castrated male Yucatan minipig may serve as a better model for male human SC tissue rather than human female SC tissue.

Finally, it is important to point out reports related to minipig lipids having a higher proportion of stearic and oleic fatty acids and less palmitic acid, as compared to human tissues exist [33, 34]. This affects the overall viscoelasticity as the fatty acid composition can alter the fluidity and packing of the lipids in the cell membrane, affecting the overall mechanical properties of the tissue [19, 35]. Stearic and oleic fatty acids are unsaturated fatty acids that increase the overall fluidity of the cell membrane, making it more flexible. This leads to a decrease in its elastic (storage) modulus, with an increase in the viscous (loss) modulus of the tissue [19, 20]. On the other hand, palmitic acid, which is a saturated fatty acid, makes the cell membrane more rigid, leading to an increase in bulk-elastic modulus and a decrease in the bulk-loss modulus of the tissue [19, 20]. Quantifications of the abovementioned fatty acids would be required in developing a more detailed understanding of the biochemical cues for the observed bulk-viscoelastic behavior.

In summary, this study is able to decouple tissue viscoelasticity and its dependence on lipid concentrations for minipig skin tissues at different body locations and depths. This study points out the significant differences that exist among the rheological behavior and biochemical composition of minipig and human SC tissue models. The minipig neck (breast) showed the highest (lowest) storage modulus across all the rheological tests. The closest agreement in storage modulus, between all three minipig locations was observed in layer 3. A similar trend was noted for the relaxation behavior. We observed that reduced strain-stiffening at high strain amplitudes is caused by higher lipid content within the tissue. Concluding the existence of, species, anatomical location, tissue depth, sex, and, strain dependent rheological properties. The findings also suggest a male Yucatan minipig might present a better model for male human SC tissue, as compared to human female SC tissue. Hence, we expect the outcomes from this study to motivate future works, in avoiding generalization of the viscoelasticity for minipig-skin tissues over human SC tissues when used for comparison purposes. Hopefully, this will stimulate more detailed comparative studies on body location and depth-based analysis, exploring both the micro/macro-mechanical properties in relation to the biochemical cues.

## Acknowledgments

This research was supported by a grant from Eli Lilly and Company. The authors would also like to thank, Jordanna M. Payne for her assistance in acquiring the tissue samples and Randy Ewoldt for providing the MITlaos software.

## A Rheological data fit parameters

The loss modulus from the frequency and amplitude sweeps are represented in Figs. 11 and 12, respectively. The results of the power law and GM model fit are presented in Tables 1 and 2, respectively. Relaxation half-life times for all the tissues are shown in Table 3.

**Fig. 11.**
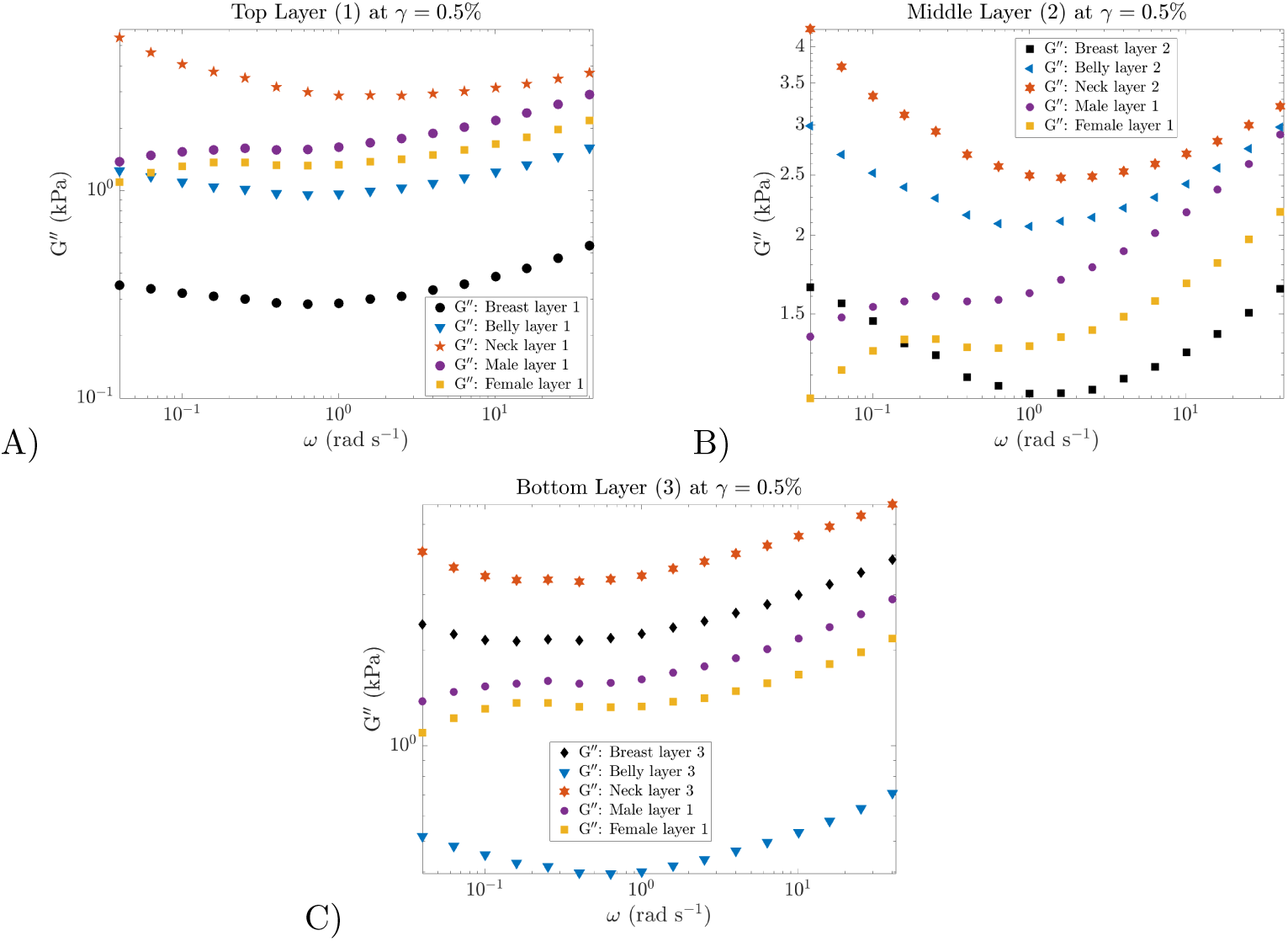
Loss modulus, G*^′′^*(*ω*), for the three different regions of the minipig, i.e., breast, belly, and neck, across the different layers, 1-3, represented through a-c, respectively. The male and female human SC samples are overlaid in all the figures for comparison.

**Fig. 12.**
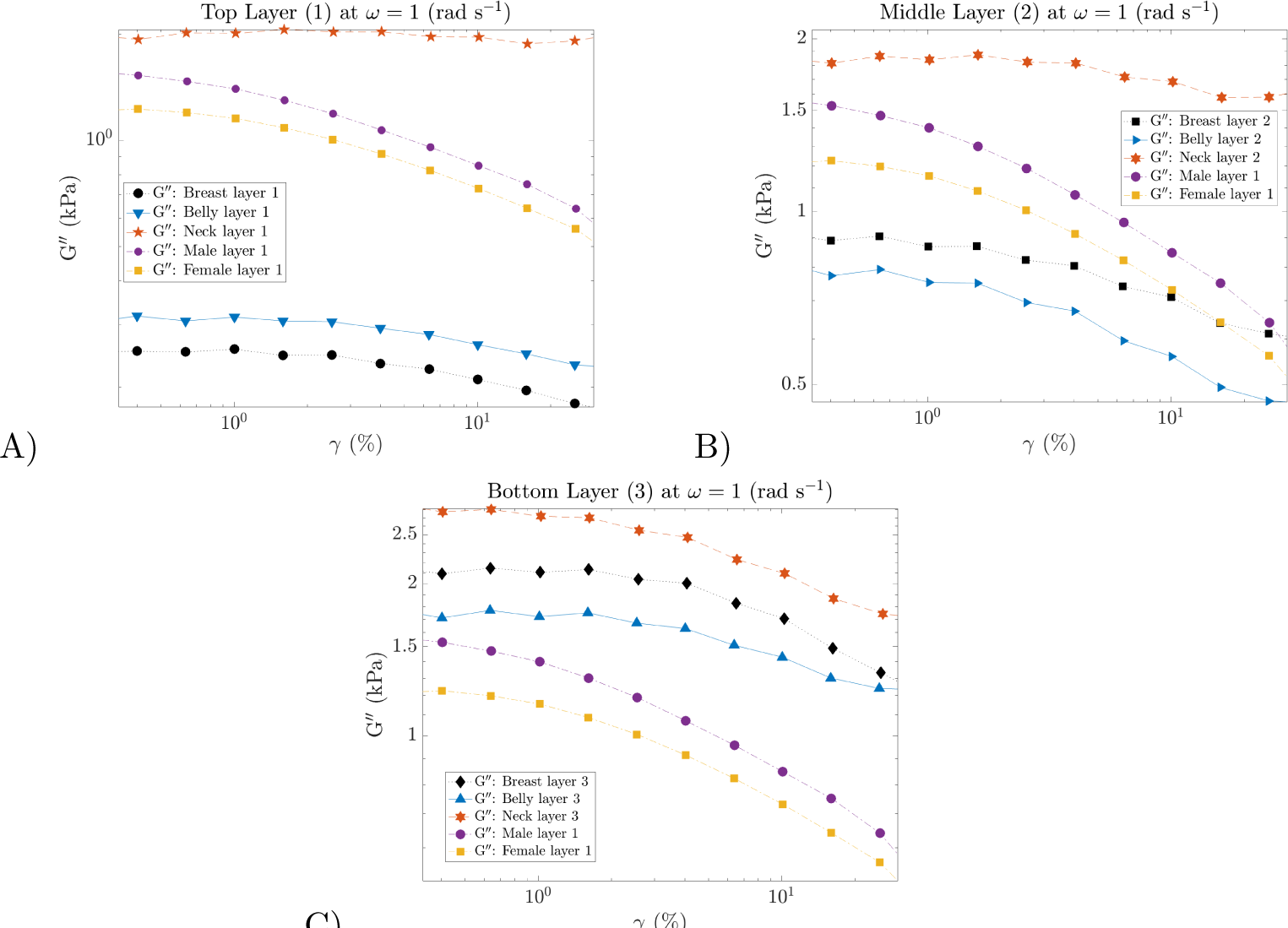
Loss modulus, G*^′′^*(*γ*), for the three different regions of the minipig, i.e., breast, belly, and neck, across the different layers, 1-3, represented through a-c, respectively. The male and female human SC samples are overlaid in all the figures for comparison. Continuous lines are overlaid in order to guide the reader and are not a fit.

## B LAOS

The degree of elastic and viscous non-linearities are quantified in Tables 4 and 5 respectively. The LB curves for the two strains, 15% and 40%, are illustrated for all the cases in Fig. 13. The increased distortions at 40% strain in Fig. 13 reveal a strong nonlinear response.

**Fig. 13.**
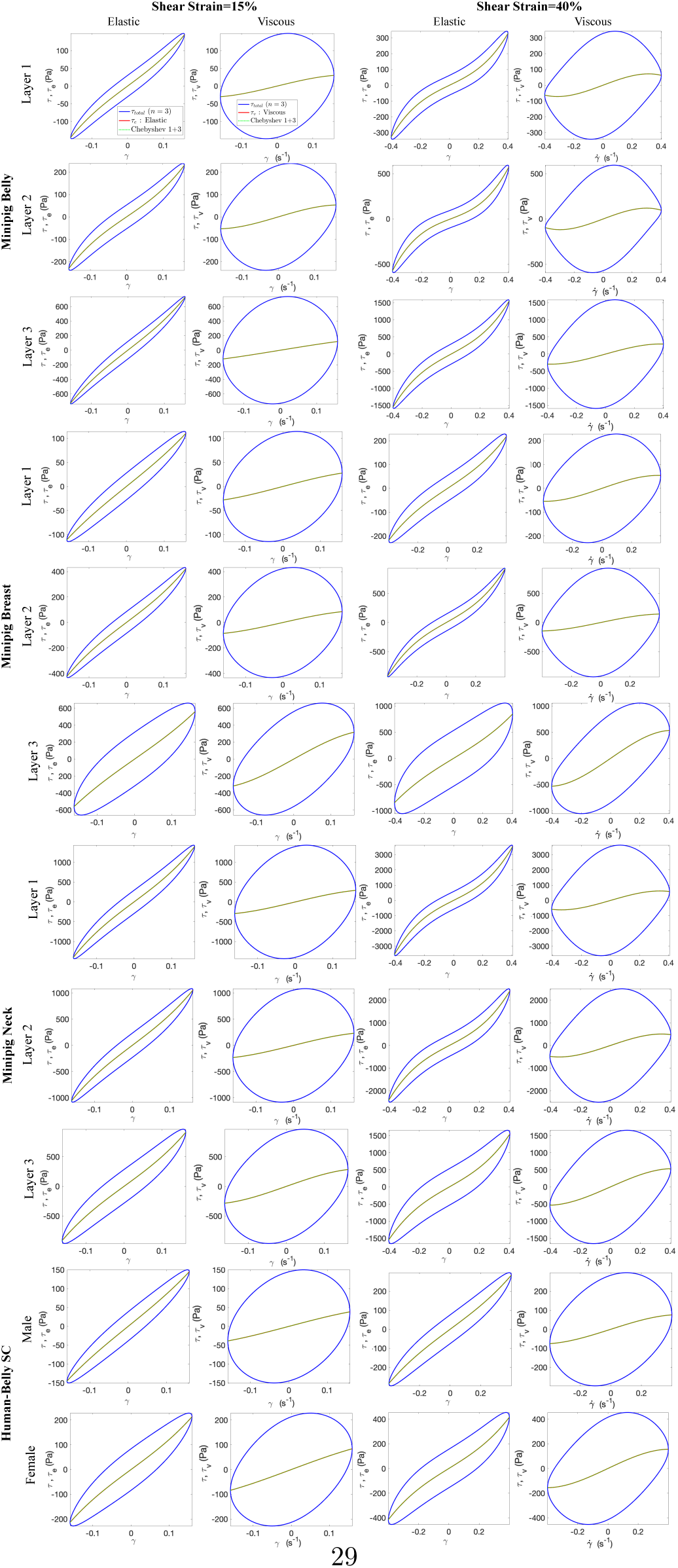
LB curves plotted using MITLaos of all the tissues at 15% and 40% shear-strain amplitudes. The blue (red) continuous curves are for the total stress (elastic/viscous) filtered for the 3*^rd^* harmonic. The area enclosed by the LB curves is related to the energy dissipated per unit volume in one complete cycle of the oscillatory strain. The legend shown in the case for the elastic behavior of the minipig-belly layer 1 at 15% strain is the same for all the figures.

